# Modulation of white matter bundle connectivity in the presence of axonal truncation pathologies

**DOI:** 10.1101/2020.01.14.903559

**Authors:** Robert E. Smith, Fernando Calamante, Sanuji Gajamange, Scott Kolbe, Alan Connelly

**Affiliations:** The Florey Institute of Neuroscience and Mental Health, Melbourne, Australia; The Florey Department of Neuroscience and Mental Health, The University of Melbourne, Melbourne, Australia; Sydney Imaging, The University of Sydney, Sydney, Australia; School of Biomedical Engineering, The University of Sydney, Sydney, Australia; Department of Neuroscience, Monash University, Melbourne, Australia

## Abstract

Endpoint-to-endpoint fibre bundle connectivity estimated using spherical deconvolution & streamlines tractography in diffusion MRI may be excessive in the presence of pathologies that involve truncation of axons within the white matter. Here we propose a simple modification to an existing method that directly quantifies and corrects for this over-estimation.

## Introduction

The Spherical-deconvolution Informed Filtering of Tractograms (SIFT) method^1^ - as well as its successor, SIFT2^2^ - aims to imbue a whole-brain streamlines tractography reconstruction with quantitative properties. This is achieved by enforcing consistency between the density of the tractogram and estimates of fibre density from the spherical deconvolution model; following such, those streamlines attributed to any particular white matter pathway of interest provide an estimate of intra-cellular cross-sectional area of that pathway.

In a recent sequence of publications^3–5^ it was noted that in instances where such fibre cross-sectional area is not preserved along the entire bundle length, the connectivity derived from SIFT and related methods may be an over-estimate (Figure 1). This may occur for instance in the case of Wallerian degeneration, where the proximal portion of a truncated axon still contributes to the diffusion-weighted signal on which fibre density estimates are based, but does not contribute biologically to the endpoint-to-endpoint connectivity of the corresponding white matter bundle. Here we propose a modification to white matter fibre bundle connection strength estimatesderived using the SIFT2 method to account for such pathologies.

**Figure 1.**
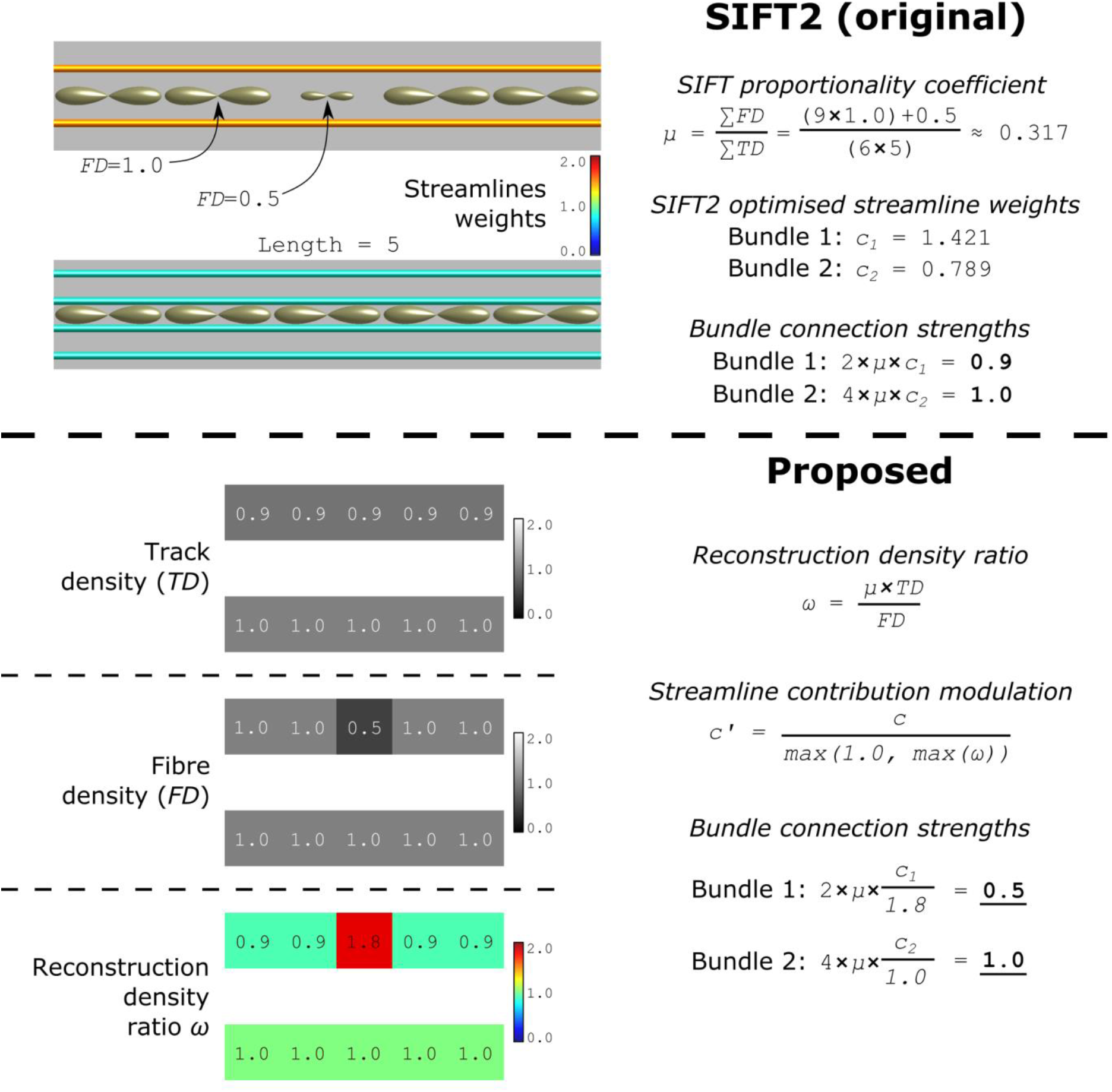
Top: The phantom structure under discussion in prior publications, and connectivity quantified by SIFT2. Bottom: Derivation and application of the proposed modulated connectivity metric.

## Methods

In SIFT2, for each individual streamline, an appropriate “weight” is calculated, such that the aggregate of all streamlines and their corresponding weights minimises the differences between streamlines and fibre densities throughout the image. These “weights” physically represent cross-sectional area multipliers, and it is the sum of these weights for those streamlines ascribed to a particular bundle that quantifies the total intra-cellular cross-sectional area - and hence “connectivity” - of that bundle.

The relationship between streamlines and fibre density in any given location in the image can be expressed as the *reconstruction density ratio ω*. Where the local streamlines density from the tractogram model fit exceeds that of the underlying fibre density, this parameter will be greater than unity, in direct proportion to the magnitude of this excess. As such, this parameter - specifically the *maximum* of this parameter along the length of each individual streamline - serves as a measure the appropriate quantity by which to scale the contribution of each streamline, preventing the reconstruction from expressing a streamlines density that is greater than the fibre density estimated by the diffusion model. Figure 1 demonstrates how this provides a measure of endpoint-to-endpoint connectivity that precisely mimics the information-carrying capacity of the synthetic pathology introduced into the phantom.

## Results

Figure 2 shows the spatial distribution of fibre density estimated by spherical deconvolution throughout an exemplar coronal slice for a patient with Multiple Sclerosis (MS), in comparison to spatial distributions of streamline densities contributing to endpoint-to-endpoint connectivity, with and without the proposed modification. SIFT2 over-estimates the density of white matter fibres in the highlighted pathological region. Applying the proposed modulation to streamlines weights across the entire white matter results in an erroneous global decrease in streamlines density, due to outlier sensitivity when taking the maximum of *ω*. By instead constraining the application of the proposed connectivity modulation based on partial volume fractions of an explicit pathological tissue segmentation (which could be derived manually or automatically), the decrease in streamlines density is limited to pathological regions and the white matter pathways passing through them.

**Figure 2.**
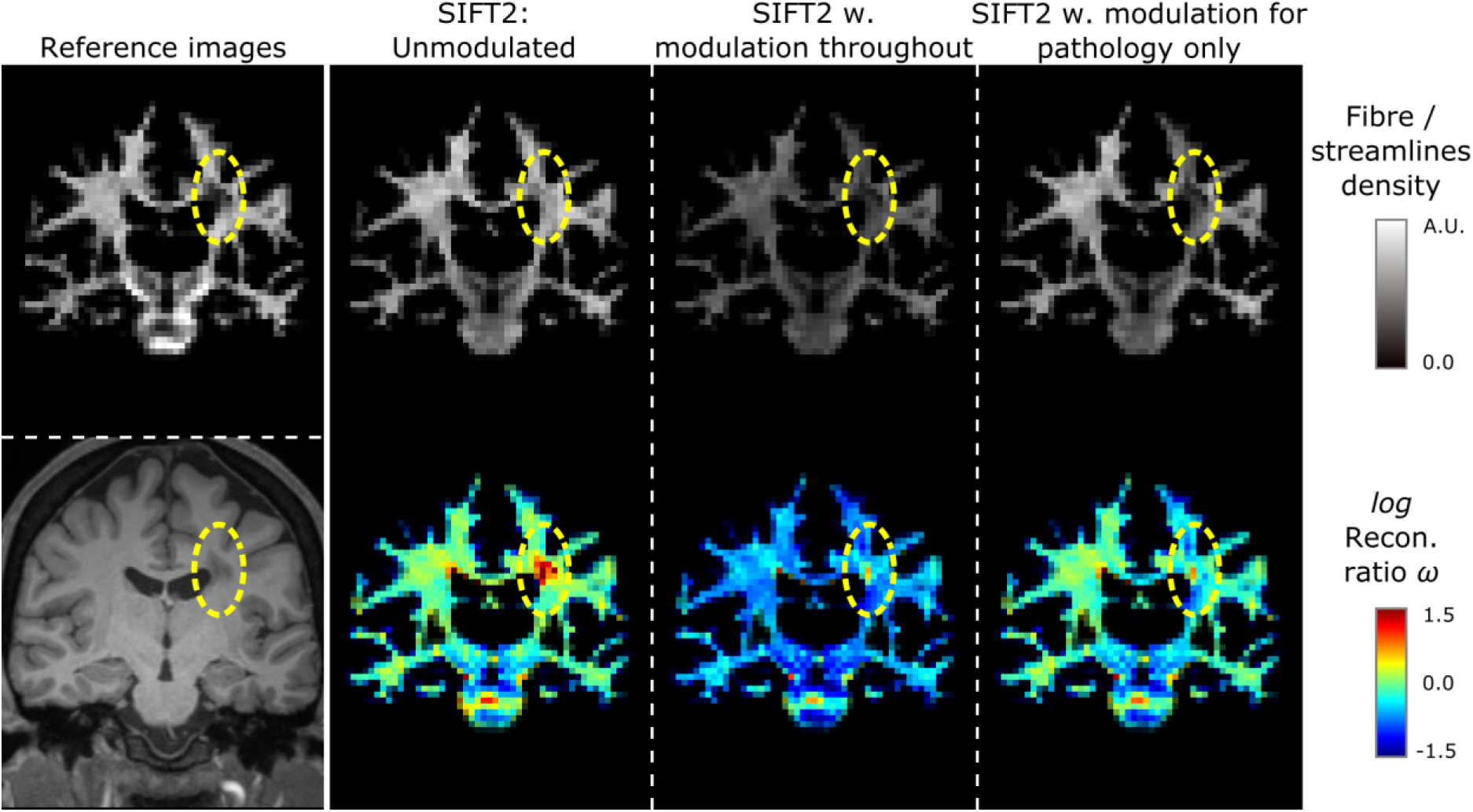
Left: Fibre density estimated through spherical deconvolution, and T1-weighted image reference. Right: Spatial distributions of streamlines density with and without proposed modification.

## Conclusions

Where endpoint-to-endpoint white matter connectivity is to be quantified in the presence of axonal truncation pathology, the proposed change to the SIFT2 method ensures that those estimates better reflect the restriction of bundle information-carrying capacity imposed by such pathology.

## Acknowledgments

We are grateful to the National Health and Medical Research Council (NHMRC) of Australia, and the Victorian Government’s Operational Infrastructure Support Program for their support. R. Smith is a fellow of the Australian National Imaging Facility, a National Collaborative Research Infrastructure Strategy (NCRIS) capability, at the Florey Institute of Neuroscience and Mental Health.

## References

1. Smith RE, Tournier J-D, Calamante F, Connelly A. SIFT: Spherical-deconvolution informed filtering of tractograms. NeuroImage. 2013;67(0):298–312.

2. Smith RE, Tournier J-D, Calamante F, Connelly A. SIFT2: Enabling dense quantitative assessment of brain white matter connectivity using streamlines tractography. NeuroImage. 2015;119:338–351.

3. Sarwar T, Ramamohanarao K, Zalesky A. Mapping connectomes with diffusion MRI: deterministic or probabilistic tractography? Magnetic Resonance in Medicine. 2019;81(2):1368–1384. doi:10.1002/mrm.27471

4. Smith RE, Calamante F, Connelly A. Mapping connectomes with diffusion MRI: Deterministic or probabilistic tractography? Magnetic Resonance in Medicine. 2020;83(3):787–790. doi:10.1002/mrm.27916

5. Zalesky A, Sarwar T, Ramamohanarao K. A cautionary note on the use of SIFT in pathological connectomes. Magnetic Resonance in Medicine. 2020;83(3):791–794. doi:10.1002/mrm.28037

